# View, engage, predict: enhancing brain-behavior mapping with naturalistic movie-watching fMRI

**DOI:** 10.1101/2025.07.28.666907

**Authors:** Yunrui Zhang, Emily S. Finn, Mert R. Sabuncu, Amy Kuceyeski

## Abstract

Most brain-behavior mapping studies rely on resting-state functional connectivity (FC), but this approach has known accuracy limits and can be outperformed by movie-watching FC. Here, we present a novel deep neural network framework to predict cognitive scores and sex from FC during naturalistic movie viewing, and examine how movie content and its ability to synchronize brain activity across individuals relate to prediction performance. We show that FC from movie-watching generally outperforms resting-state FC - even when compared to five times more temporal data - with sensory and higher-order brain networks emerging as the most important for prediction. Using both static and sliding-window dynamic FC approaches, we find that higher cognitive prediction accuracy is significantly associated with greater inter-subject synchrony and the duration of human faces and voices in the movies; these effects were not found for sex prediction. This work underscores the promise of naturalistic movie viewing as a powerful tool for probing individual differences in the brain and revealing neural underpinnings of human behavior.

## 1. Introduction

Understanding brain regions and circuits that underlie individual differences in cognition and behavior is a fundamental goal in neuroscience. Functional MRI, which allows non-invasive recording of brain activity, has revealed that functional connectivity (FC) patterns are uniquely identifiable across individuals, i.e. FC fingerprinting, and are associated with various behavioral traits [1] [2] [3] [4] [5]. Traditionally, FC fingerprinting and brain-behavior mapping has relied on resting-state fMRI, but naturalistic paradigms such as movie watching offer a more ecologically valid and engaging framework for that at times results in more accurate brain-behavior mappings [6] [7] [8] [9] [10]. Watching a movie induces rich and dynamic neural responses, characterized by synchronized activity across individuals and structured temporal fluctuations in functional connectivity [11] [12] [13] [14]. The use of movie stimuli provides a controlled yet immersive setting, capturing brain activity fluctuations that more closely resemble real-world cognitive processes compared to static resting-state paradigms, in which a person is told not to think about anything in particular.

Recent studies have used FC to predict behavior, i.e. cognitive scores, motor outcomes or diagnosis categories, or demographics like age or sex, but existing models and data remain limited in their predictive power. Most prior brain-behavior mapping studies have employed linear models like linear regression [1] [15]; however, these methods may be limited, as the mapping between functional connectivity (FC) and behavior may involve nonlinear components. The application of deep learning to FC-based behavior prediction has been an area of recent exploration, and may have the potential to uncover complex mappings that traditional models overlook [16] [17] [18] [19] [20] [21]. In this study, we introduce a novel deep neural network (DNN) framework for predicting cognition and sex from FC patterns derived from movie-watching fMRI data, leveraging machine learning techniques to enhance predictive performance and interpretability. One commonly cited drawback of DNNs is that they suffer from a lack of explainability, which is an important aspect of brain-behavior mapping. Being able to reliably identify the functional connections, regions or networks that are important for predicting a given task are of utmost importance for the field of network neuroscience [22] [4] [23] [24] [25]. Knowing the neural underpinnings of behavior or diagnoses would enable better understanding of how we may intervene to enhance performance or reduce symptoms.

Another largely unexplored aspect of FC-based behavior prediction using movie-watchin data is its relationship with neural synchrony, or the level of temporal alignment across individuals during shared experiences [26] [27] [28] [29] [30]. Increased neural synchrony has been associated with higher levels of attentional engagement and cognitive processing [31] [32]. However, the connection between neural synchrony and brain-behavior prediction accuracy remains under explored.

Finally, prior research has largely focused on static functional connectivity, overlooking the temporal dynamics of FC during movie-watching. While static FC provides an aggregate measure of connectivity over an entire scan or movie clip, it cannot capture the moment-to-moment fluctuations that occur in response to evolving stimuli [33] [34]. Dynamic functional connectivity (dFC) can reveal transient connectivity states that may be more informative for predicting behavior [35] [36] [37] [38] [39] [40], and in the context of brain-behavior mapping allow finer scale analysis of the movie features that may be driving prediction accuracy.

In this study, we introduce a novel deep learning-based framework for predicting cognition and sex from functional connectivity during movie-watching and at rest. We compare the predictive performance of various movie clips with differing content, systematically analyzing the contributions of individual brain connections and regions to the predictions. Additionally, we explore the relationship between neural synchrony and brain-behavior prediction accuracy to test our hypothesis that higher neural synchrony generally correlates with better prediction performance. At a finer temporal scale, we segment the movie clips into sliding windows to capture the dynamic fluctuations in neural synchrony and prediction accuracy, further testing for correlations between synchrony and prediction performance. Finally, we investigate the visual, auditory, and semantic features of the movies, identifying key factors that correlate with predictive performance at both the level of the entire movie clip as well as the sliding windows within a clip. By integrating these approaches, our work provides novel insights into the relationship between functional brain connectivity, behavior, neural synchrony, and the features of naturalistic stimuli, offering a more comprehensive understanding of the brain and stimulus features that drive brain-behavior mapping using naturalistic paradigms.

## 2. Results

### 2.1 Brain-behavior mapping accuracy varies across movie clips

We applied a deep neural network (DNN) to the static FC extracted from different movie clips (and rest) to predict individual traits, including cognitive scores and sex, in a cohort of 176 participants (106 female, 70 male, aged 22-36) from the 7T substudy in the Human Connectome Project’s Young Adult dataset [41]. Cross-validation was performed with 10-folds, ensuring family members did not span folds; this cross-validation was repeated 100 times to assess robustness. The prediction accuracy for cognitive scores was quantified using the Spearman correlation coefficient between true and predicted scores across participants in the test set, then averaged across the 100 iterations. To assess statistical significance, p-values were computed by randomly permuting the predicted scores 10,000 times in each iteration. We report the median p-value across all 100 iterations. For sex classification, prediction performance was evaluated using the area under the receiver operating characteristic curve (AUC) for the test set, also averaged over 100 iterations. Our results show wide variability in the predictive performance across the different movie clips. Most movie-based FCs outperformed the rest FC calculated using the same duration of fMRI data, whereas using the full 15 mins of resting FC gave accuracy that was higher than most movie clip FC models. However, the best performing movie clip FC models, each with only 3 mins of fMRI data, outperformed resting state FC models using 15 mins of data (Fig. 2a,b).

**Figure 1:**
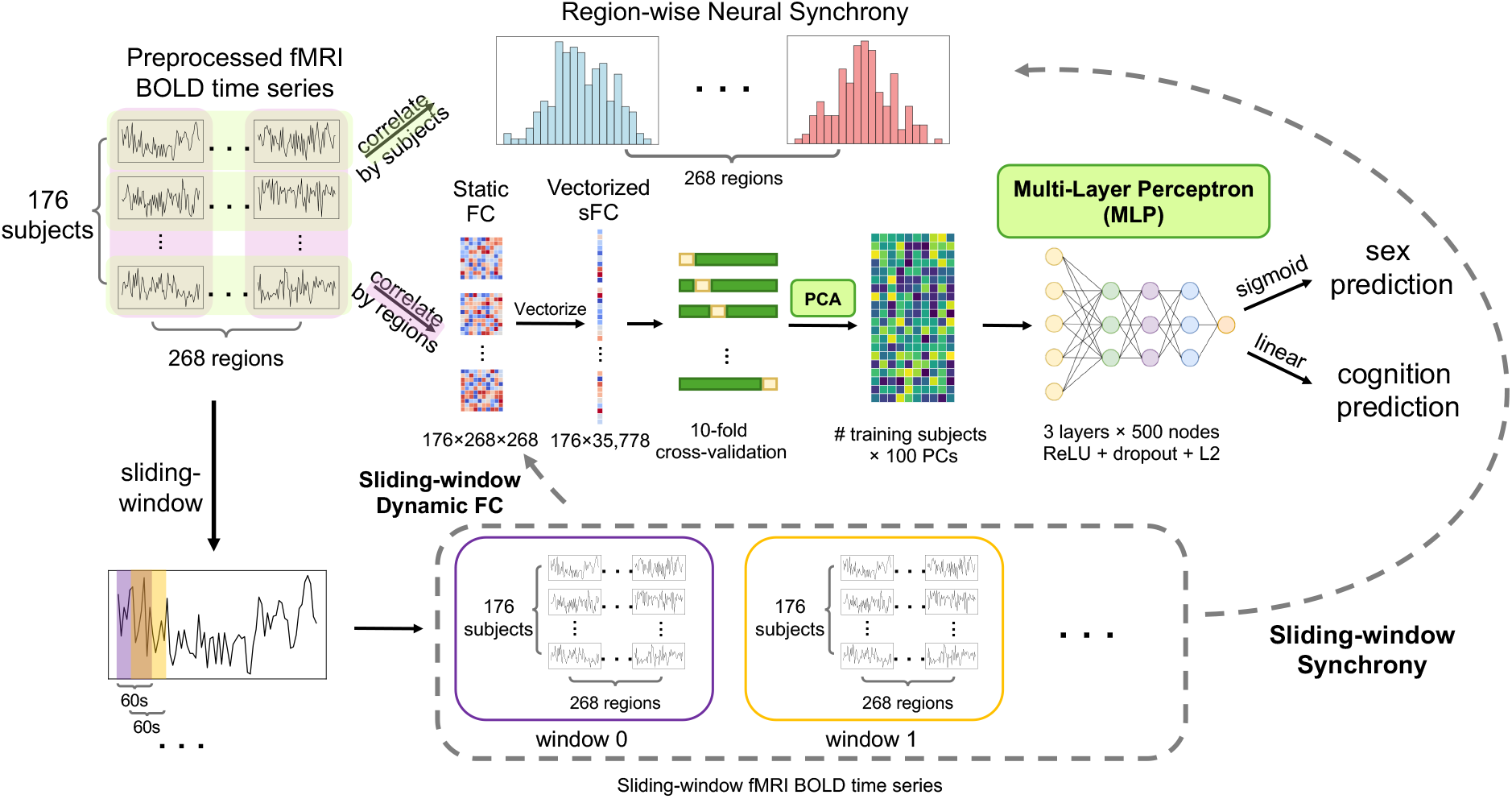
Workflow of the study. Static functional connectivity (sFC) is computed by correlating fMRI time series across pairs of brain regions, followed by vectorization and PCA to reduce dimensionality. Neural synchrony is defined by across-subject correlations of the same brain regions’ fMRI time series and assessed by the mean/std of distribution across pairs of subjects. In parallel, a sliding-window approach is applied to compute dynamic functional connectivity (dFC) and sliding-window neural synchrony, capturing temporal fluctuations in connectivity patterns. The extracted features from both static and dynamic FC are fed into a 3-layer multi-layer perceptron (MLP) model with 500 nodes per layer for classification (sex prediction) and regression (cognition prediction).

**Figure 2:**
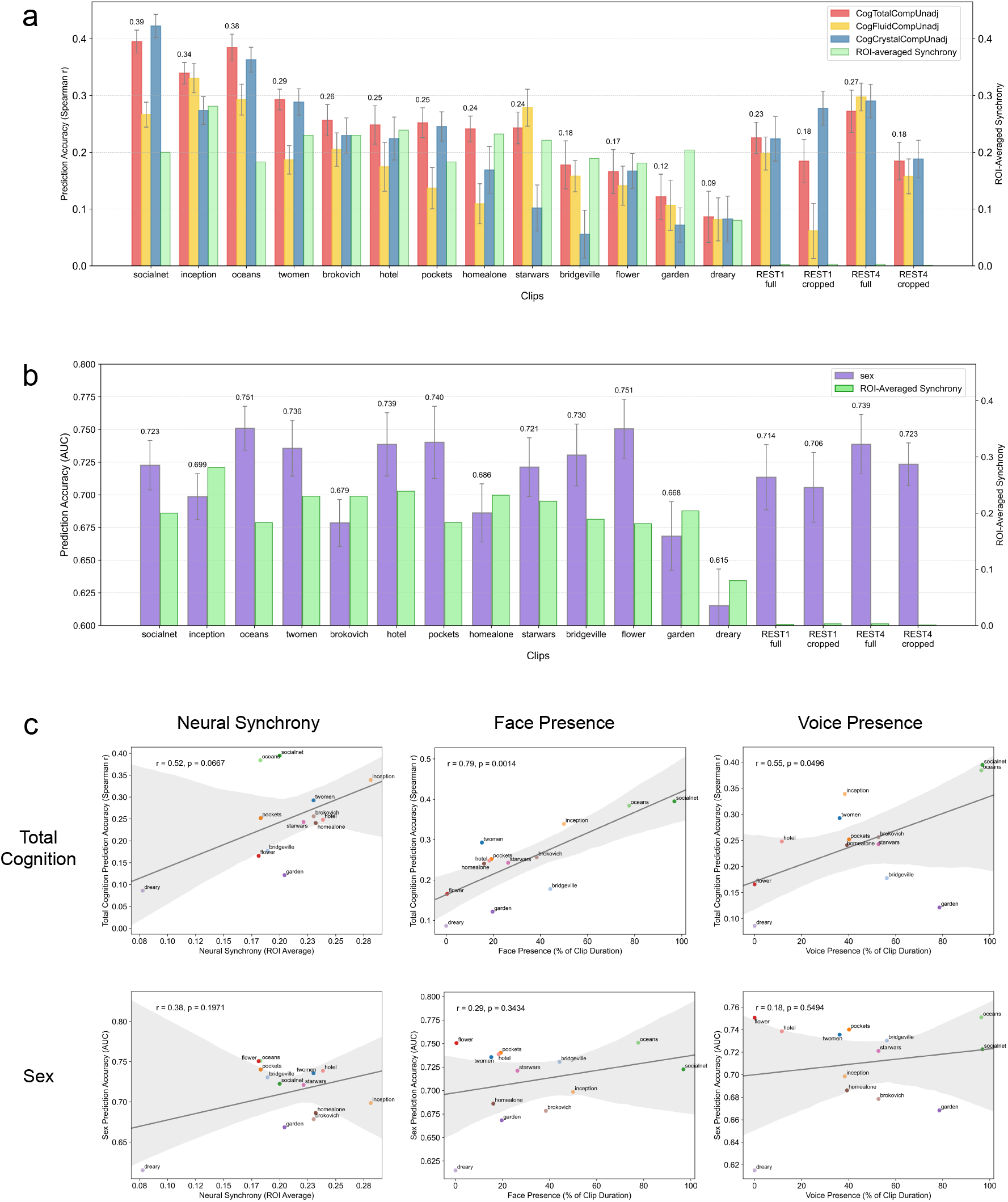
Brain-behavior mapping model performance. (a) Prediction accuracies for total, fluid, and crystallized composite cognition scores across all movie clips and resting-state sessions (both full and cropped), evaluated using Spearman’s rank correlation coefficient (*r*). The colored bar height is the average and the grey lines are the standard deviation of the accuracy values over the 100 cross-validation iterations. All sessions were cropped to 164 seconds from onset to ensure same input length across sessions, except for the full resting-state fMRI scan that consists of 15 mins of data. Regional average neural synchrony, over 55 brain regions with significant synchrony in at least one movie clip, averaged across all pairs of subjects are shown in green. (b) Prediction accuracies for sex classification across all movie clips and resting-state sessions, assessed using area under the curve (AUC); bar height is the average value and the grey lines are the standard deviation of the accuracies over the 100 iterations. Regional average neural synchrony over all pairs of subjects are shown in green. (c) Across movie clips, neural synchrony is positively and significantly correlated with total cognition prediction accuracy (*r* = 0.52, *p* = 0.0667) and positively but not significantly related to sex classification accuracy (*r* = 0.38, *p* = 0.1971). Total cognition prediction accuracy is significantly correlated with the percentage of clip duration containing human faces (*r* = 0.79, *p* = 0.0014) and voices (*r* = 0.55, *p* = 0.0496). Sex prediction accuracy also shows positive, though weak and non-significant, correlations with face presence (*r* = 0.29, *p* = 0.3434) and voice (*r* = 0.18, *p* = 0.5494) presence.

For total cognition, the highest prediction accuracies were observed for models based on movie-watching FC, including *Social Network* (Spearman *r* = 0.39 ± 0.02, permutaion-based *p* = 0), *Ocean’s 11* (*r* = 0.38±0.02, *p* = 0), and *Inception* (*r* = 0.34±0.02, *p* = 0). In contrast, the lowest predictive performance was associated with clips from the third movie session, including *Flower* (*r* = 0.17±0.04, *p* = 0.043), *Garden* (*r* = 0.12±0.04, *p* = 0.078), and *Dreary* (*r* = 0.09±0.05, *p* = 0.120). The two rest sessions also demonstrate predictive power, with *REST1* (*r* = 0.18 ± 0.04, *p* = 0.034) corresponding to the same day as the first and second movie sessions, and *REST4* (*r* = 0.18 ± 0.03, *p* = 0.033) aligning with the third and fourth sessions (Fig. 2a). Note that all movie clips and rest sessions were cropped to 164 seconds from their onset to ensure comparability across conditions. The 15-min full rest sessions show better prediction accuracy than the cropped rest sessions (full *REST1* : *r* = 0.23 ± 0.02, *p* = 0.009; full *REST4* : *r* = 0.27 ± 0.03, *p* = 0.002), but this result is outperformed by the movie clips from *Social Network, Inception, Ocean’s 11* and *Two Men*, which are 5 times shorter than the full rest sessions. The relative ranking of prediction accuracy for other cognitive measures, including crystallized cognition and fluid cognition, followed a similar trend to that of total cognition (Fig. 2a). However, there are slight variations in the prediction accuracy ranking for crystallized and fluid cognition; for example, while *Social Network* yielded the highest predictive accuracy for total cognition (*r* = 0.39 ± 0.02, *p* = 0) and crystallized cognition (*r* = 0.42 ± 0.02, *p* = 0), *Inception* exhibited the highest predictive accuracy for fluid cognition (*r* = 0.33 ± 0.03, *p* = 0). Results for all 12 individual NIH toolbox cognitive measures and chronological age are presented in Supplementary Figure 1; in the main text, we do not present the individual cognitive scores as they are noisier than the composites and we do not present age as the spread of age values is quite small in this population and thus the accuracy quite low.

For sex classification, the highest predictive accuracies were observed for *Ocean’s 11* (*AUC* = 0.75 ± 0.02), *Flower* (0.75 ± 0.02), and *Pockets* (0.74 ± 0.03), whereas *Erin Brockovich* (0.68 ± 0.02), *Garden* (0.67 ± 0.03), and *Dreary* (0.62 ± 0.03) demonstrated the lowest predictive power (Fig. 2b). 164-second *REST1* (0.71 ± 0.03) and *REST4* (0.72 ± 0.02) demonstrated predictive accuracies that were either lower or higher than those of specific movie clips, as observed in cognition prediction. The 15-min full rest sessions show slightly better prediction accuracy (full *REST1* : *AUC* = 0.71 ± 0.02, full *REST4* : 0.74 ± 0.02), which is outperformed by shorter movie clips from *Ocean’s 11, Flower, Pockets* and *Hotel*. Notably, while some movie clips consistently outperformed rest sessions in both cognition and sex prediction, the ranking of prediction accuracies for sex classification differed from that of total cognition.

### 2.2 Key Brain Connections and Networks for Multiple Cognitive Measures and Sex Prediction Partially Overlap in Movie-Watching Sessions

We evaluated the contribution of specific brain connections and functional networks to cognition and sex prediction using the Integrated Gradients (IG) method. The importance of individual connections was quantified by calculating the absolute IG values derived from the deep neural network (DNN) model, averaged across all 176 subjects. We identified the connections with the top 1% average importance scores and mapped to their corresponding Yeo 7 functional networks [42], and visualized using chord diagrams(Fig. 3 and, alternatively, glass brain visualizations in Supplementary Figure 3). During movie-watching sessions, the most predictive connections involved interactions between sensory networks—including the visual (VIS) and auditory (within the somatomotor network, SMN)—and high-order networks such as the default mode network (DMN), frontoparietal network (FPN), dorsal attention network (DAN), and salience network (SAN). During the two rest sessions, DMN, FPN and SAN are the most important for behavior prediction. Although the most predictive connections across movie clips often originate from the same functional networks, the clip-specific top connections vary considerably. For example, in *Inception*, the most predictive connections were primarily within high-order networks (DAN and DMN), whereas in *Ocean’s 11*, they were more concentrated within the lower-order visual network. Notably, during both movie-watching and rest sessions, many connections important for cognition prediction also played a significant role in sex classification. The overlap between cognition and sex prediction was more pronounced within the same clip than across different clips. This indicates that perhaps the most important functional connections underlying behavior and demographic prediction may be primarily modulated by the features of the movie stimuli.

**Figure 3:**
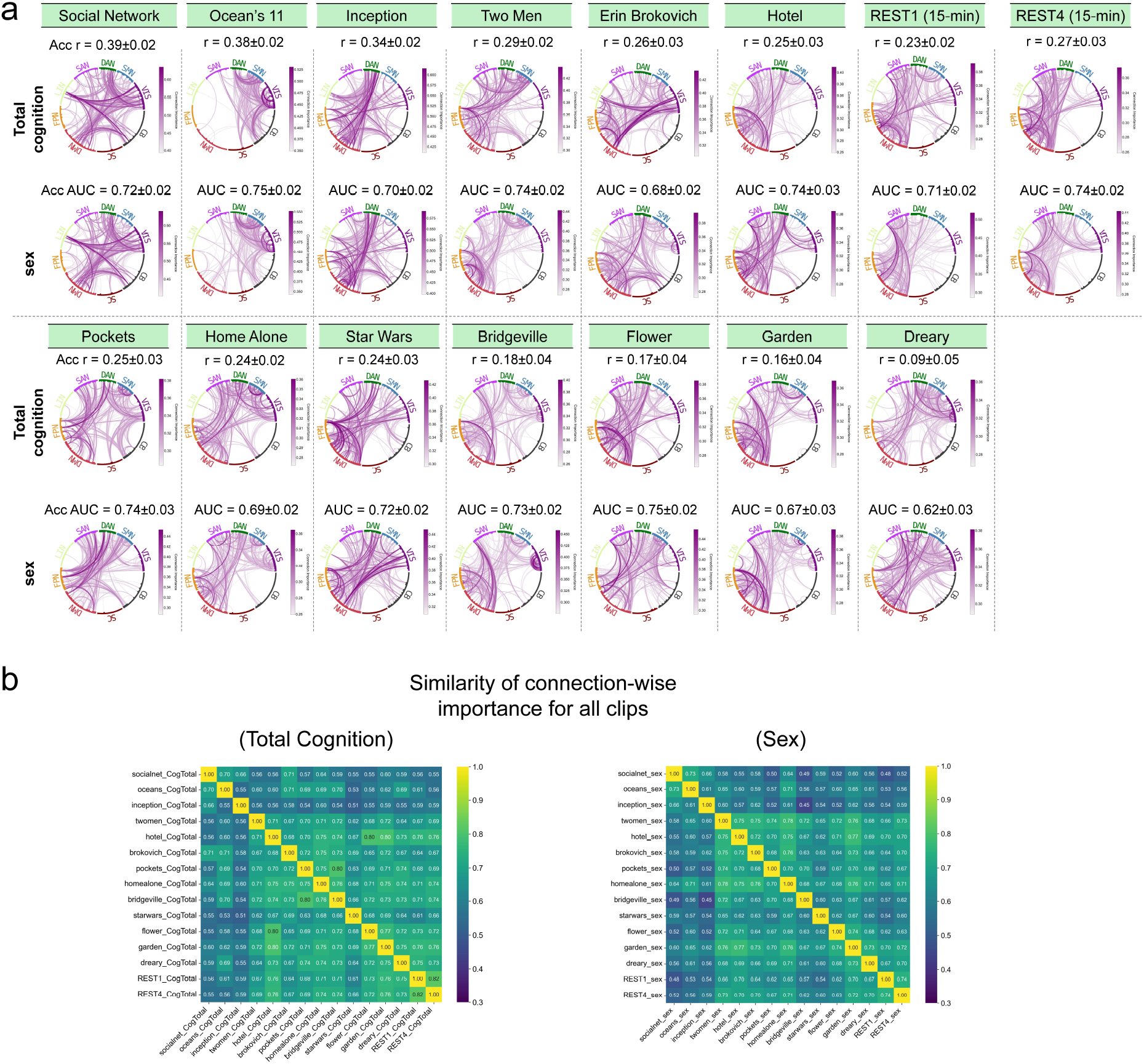
Brain connections important to total cognition and sex predictions. (a) For each clip and resting-state session, chord diagrams visualize the most important connections—specifically, those within the top 1% of average importance scores—for both total cognition and sex prediction models. The most important connections appear to vary across clips and remain consistent for total cognition and sex within a clip. These critical connections are predominantly located in sensory networks (VIS, SMN) and high-order networks (DMN, FPN, SAN, DAN), though the specific top connections vary across different movie clips. (b) Correlation of edge-wise FC importance across all clips, for total cognition prediction and sex prediction. Edge-wise FC importance similarity is assessed by correlating the importance scores of all pairwise FCs across two fMRI sessions.

To obtain a more quantitative and comprehensive analysis of the association between varied movie clips’ FC feature importances, we correlated the FC edge-wise feature importance maps for both total cognition and sex, see Fig. 3b. For the most part, the correlations ranged from moderate to high with *Ocean’s 11, Inception* and *Social Network* clips appearing somewhat as outliers in both cognition and sex predictions. We performed the same analyses across all clips and all individual NIH toolbox cognition scores (plus sex) in Supplementary Figure 4. We observe the same overall pattern of edge-wise FC feature importances being more similar within the same movie clip for different cognitive scores or sex, compared to the same cognitive score or sex across different movie clips. It appears that the clip features are driving the importance of the FCs in predicting varied cognitive outcomes and sex.

### 2.3 Neural Synchrony and Functional Connectivity-based prediction accuracy are correlated

We observe that different movie clips elicit varying degrees of regional synchrony in the fMRI BOLD responses across subjects. Intuitively, movies that induce higher neural synchrony across the brain may be more engaging for most participants, as higher engagement likely means more time-locked, stimulus-driven brain activity. In addition, as neural activity increases in stimulus-driven regions during such movies, the signal-to-noise ratio (SNR) of the BOLD response and functional connectivity likely improves, potentially enhancing behavior prediction. However, very high levels of neural synchronization could mean reduced inter-individual variability, which may limit prediction capabilities across the population. This raises the question: how does neural synchrony impact brain-behavior prediction?

To investigate this, we first examined whether neural synchrony across all movie clips was correlated with prediction accuracy (Fig. 2c). For each brain region, neural synchrony was quantified using inter-subject correlation (ISC) of preprocessed fMRI BOLD signals across the duration of a movie clip or rest session, averaged over all subject pairs. The overall synchrony for a movie clip or rest session was then computed by averaging regional synchrony values across all 55 regions that had significantly synchronized activity in at least one movie clip. We found a positive, near significant correlation between neural synchrony and total cognition prediction accuracy (Pearson *r* = 0.52, *p* = 0.0667), and a weaker, non-significant correlation between synchrony and sex prediction accuracy (*r* = 0.38, *p* = 0.1971). Although the relationships did not reach statistical significance (*p <* 0.05), the positive trend suggests that higher synchrony may contribute to improved total cognition prediction accuracy.

Regional synchrony maps showing the average inter-subject correlation (ISC) across all pairs of individuals for each movie clip and rest session are shown in Fig. 4b, top row. During most movie-watching sessions, the highest synchrony was observed in auditory, visual, and language-related regions. For example, peak synchrony reached around 0.65 in the auditory cortex during *Inception* and 0.48 in *Ocean’s 11*. Generally, auditory region synchrony was higher than visual region synchrony, which has been shown in the past[43] and may be ascribed to more variability in gaze location during the movie watching than in what the person is hearing, the latter of which should be the identical across participants. As expected, neural synchrony during rest sessions was minimal across all regions.

**Figure 4:**
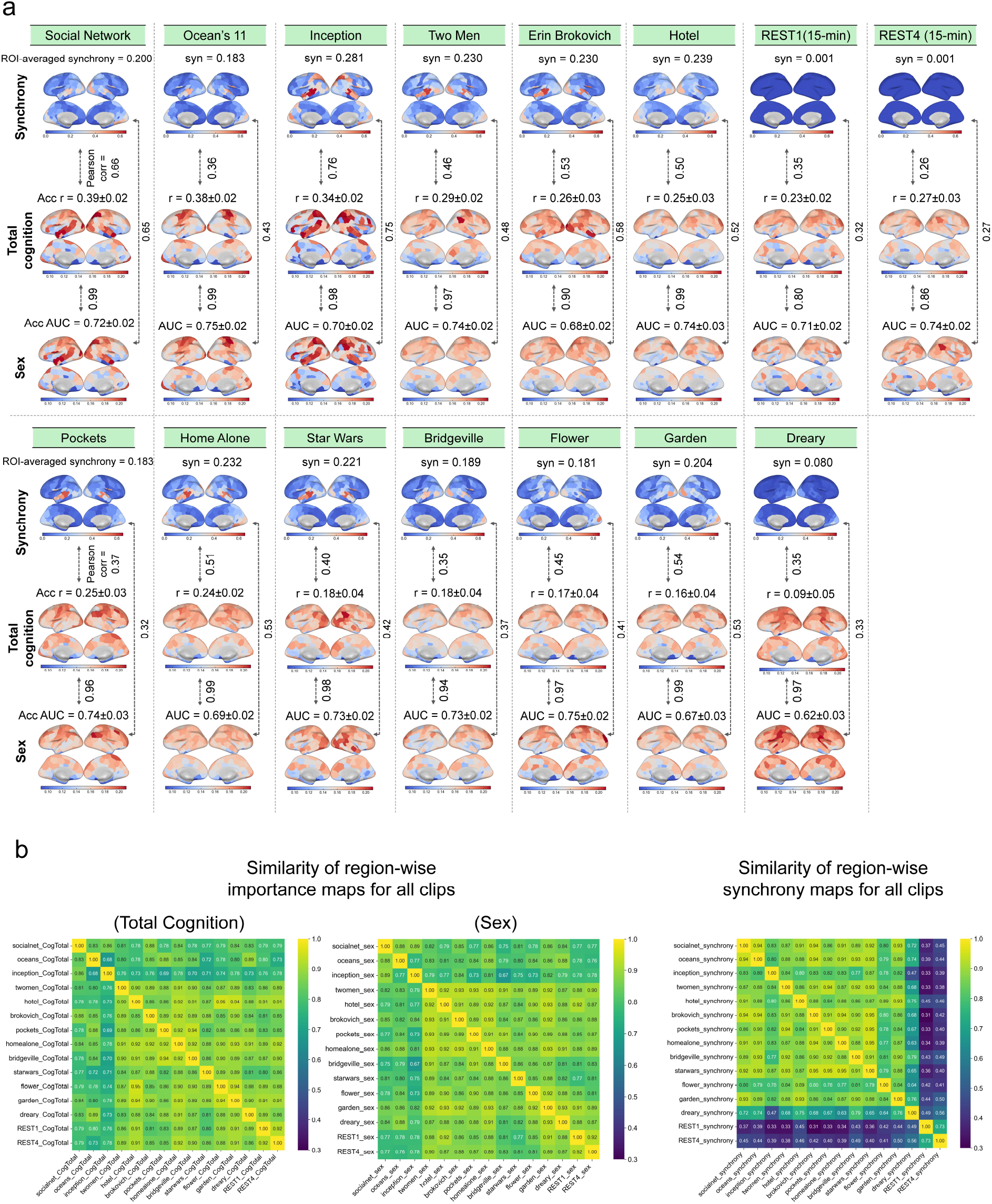
Neural synchrony and static FC feature importances. (a) For each clip, the first row shows regional neural synchrony maps (average across synchronized regions listed at the top of the panel). The second and third rows display region-wise prediction importance maps, i.e. the average of the importance scores of all edge-wise FCs to that region, for total cognition and sex, with the average prediction accuracy at the top of the panel. For all clips, neural synchrony and feature importance maps are positively correlated, indicating that regions with higher synchrony also tend to play a key role in behavior prediction. (b) Similarity of region-wise importance maps across all clips, for total cognition prediction, sex prediction and neural synchrony respectively. Consistent with results from edge-level (see Fig. 3(b)), the region-level prediction importance is positively correlated across all clips. Neural synchrony is also consistent across clips, with the exception of clips showing particularly low synchrony, such as *Dreary*, and the two rest sessions.

Next, we show region-wise feature importance maps, computed by averaging the prediction importance scores of all 267 brain connections involving that region, for total cognition and sex prediction, see Fig. 4a (middle and bottom rows). We then assessed the correlation between regional synchrony and regional feature importance for each movie clip. The correlation between cognition and sex feature importance maps for models based on the FC from the same movie clip exceeded 0.90, highlighting a strong overlap in the predictive features for both outcomes. Additionally, the synchrony maps showed consistent, moderate, positive correlations with both models’ feature importance maps, indicating that regions exhibiting higher inter-subject synchrony also tended to have greater predictive relevance for cognition and sex. See scatter plots showing these correlations in Supplementary Figure 5. Notably, we also observed considerable similarity in regional synchrony and feature importance maps across different clips, with correlations ranging from 0.67-0.95, see in Fig. 4b.

To directly evaluate the role of high-synchrony regions in behavior prediction, we conducted targeted predictions using FC in subsets of brain regions with different levels of synchrony. As shown in Supplementary Figure 6, models based on FC in regions exhibiting high synchrony consistently outperformed models based on FC of the same number of randomly selected regions in multiple movie clips, which in turn outperformed models based on FC of low-synchrony regions. These trends were consistent for both cognition and sex prediction. This finding highlights the importance of high-synchrony regions in accurate brain-behavior models using movie-watching FC.

### 2.4 Sliding-window FC prediction accuracy, Neural Synchrony and Movie Features are correlated

At a finer temporal resolution, movie clips elicit rich dynamic fluctuations in brain activity as visual and auditory stimuli unfold over time. To capture these moment-to-moment changes, we segmented each movie clip into overlapping sliding windows (window size = 60s; step size = 1s) and tracked the temporal evolution of dynamic functional connectivity (dFC) prediction accuracy, neural synchrony, and high-level movie features within each window (Fig. 5a). Several clips exhibited significant positive correlations between dynamic synchrony and total cognition prediction accuracy, including Ocean’s 11 (*r* = 0.48, permutation-based *p* = 0), Inception (*r* = 0.37, *p* = 0), Erin Brockovich (*r* = 0.26, *p* = 0), Home Alone (*r* = 0.29, *p* = 0), Bridgeville (*r* = 0.31, *p* = 0), Flower (*r* = 0.33, *p* = 0), and Garden (*r* = 0.40, *p* = 0). Similarly, significant positive correlations between synchrony and sex prediction accuracy were observed in Two Men (*r* = 0.19, *p* = 0.02), Home Alone (*r* = 0.40, *p* = 0), Bridgeville (*r* = 0.48, *p* = 0), and Flower (*r* = 0.72, *p* = 0).

**Figure 5:**
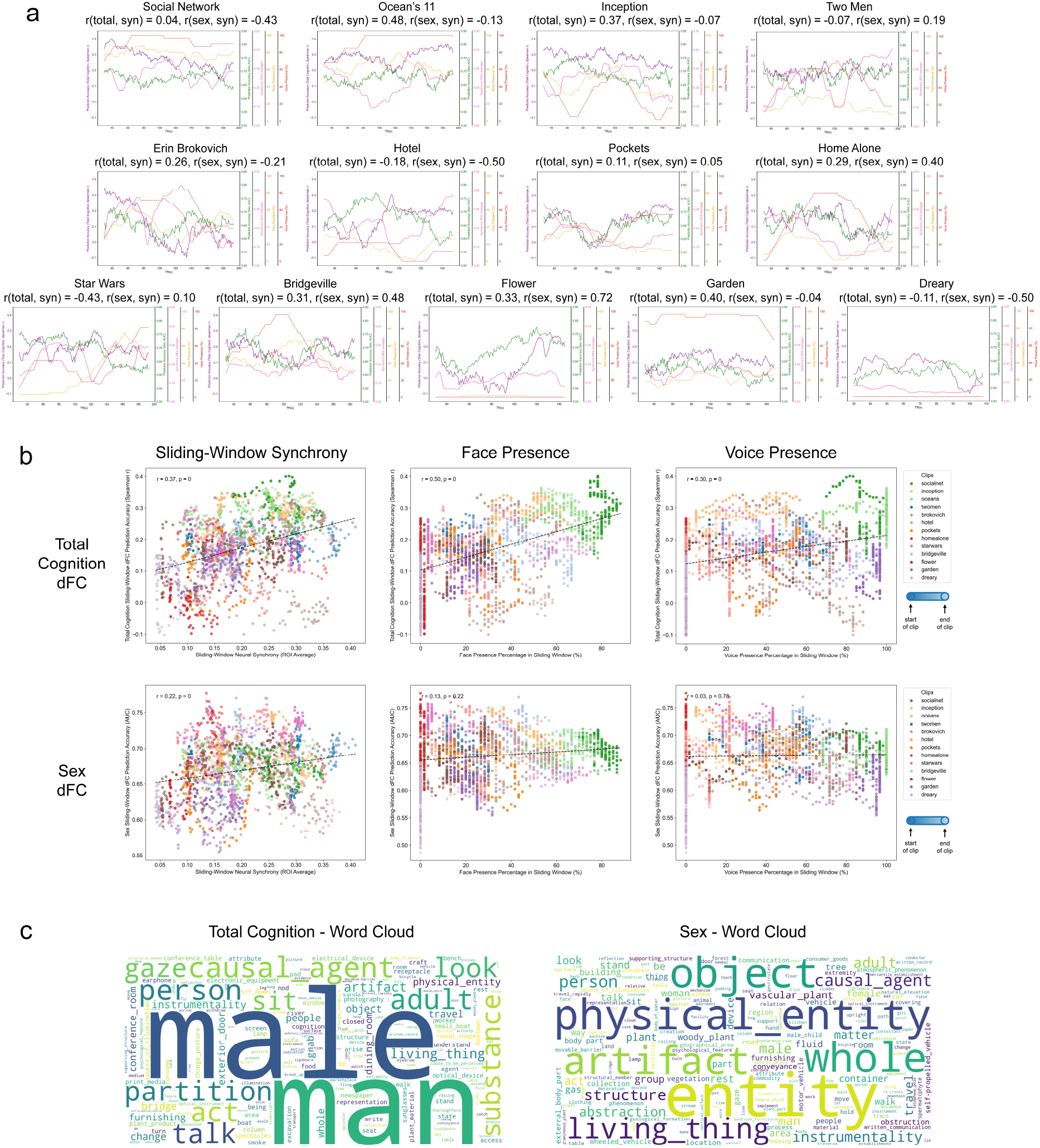
Sliding-window dynamic FC prediction, neural synchrony, and movie features co-fluctuate over time. (a) Sliding-window dynamic FC prediction accuracies for total cognition (purple) and sex (green), neural synchrony (pink), and percent of human face/voice presence (yellow/red,respectively) for all movie clips. (b) Correlations between sliding-window dynamic functional connectivity (dFC) prediction accuracies and sliding-window neural synchrony, face presence, and voice presence are computed across all sliding windows in all movie clips. Each point represents a sliding window, each color a corresponding movie clip. Point transparency indicates temporal position within the clip, ranging from 100% opacity at the beginning to 50% at the end. Sliding-window synchrony shows a positive correlation with both total cognition dFC prediction (*r* = 0.37, *p* = 0) and sex dFC prediction (*r* = 0.22, *p* = 0), with r-values calculated by Pearson correlation and p-values estimated using a 10,000-iteration permutation test. Total cognition dFC prediction is also positively correlated with face (*r* = 0.50, *p* = 0) and voice (*r* = 0.30, *p* = 0) presence percentage within sliding-windows. Sex dFC prediction was non-significantly positively correlated with face presence (*r* = 0.13, *p* = 0.22) but not correlated with voice presence (*r* = 0.03, *p* = 0.78). (c) The average frequency of WordNet features within each sliding window, weighted by that window’s dFC prediction accuracy for total cognition (left panel) or sex (right panel). The size of the words represents the weighted average count; larger words correspond to a feature that is present during more predictive sliding windows.

To gain a more comprehensive view across the different clips, we aggregated all sliding windows from all movie clips and recalculated the correlations. In this combined analysis, each point in Fig.5b represents a sliding window, colored by its corresponding movie clip. Across all windows, neural synchrony was significantly positively correlated with both total cognition dFC accuracy (Pearson *r* = 0.37, permutation-based *p* = 0) and, to a lesser extent, sex dFC accuracy (*r* = 0.22, *p* = 0). These findings are consistent with the clip-level correlations shown in Fig.2c, reinforcing the observation that periods of heightened neural synchrony are generally associated with improved cognitive and (lesser so) sex prediction.

We further explored how high-level, sliding window stimulus features modulate dynamic FC prediction performance. Total cognition dFC accuracy was positively correlated with the presence of human faces (*r* = 0.50, *p* = 0) and to a lesser extent voices (*r* = 0.30, *p* = 0). Here, face/voice presence was defined as the percentage of time points in which these features appeared cross each 60-second window. Sex prediction accuracy was not correlated with face presence (*r* = 0.13, *p* = 0.22) or voice presence (*r* = 0.03, *p* = 0.78). These results align with prior findings from static FC prediction, where the presence of human faces was associated with improved cognitive prediction accuracy [6]. Together, these results suggest that socially rich moments in naturalistic stimuli may enhance both neural synchrony and the predictive power of functional connectivity patterns in mapping to cognitive scores - but not in sex classifications.

To further investigate the semantic content underlying accurate dFC predictions, we analyzed frame-level WordNet features of the movie clips. Within each sliding window, we computed the frequency of WordNet features and weighted them by dFC prediction accuracy (for either cognition or sex). We then visualized the weighted feature distributions as word clouds (Fig. 5c), where word size reflects the weighted average frequency (weighted by prediction accuracy so more predictive windows are given larger weights) and averaged across all windows and clips. For total cognition, human- and social-related features emerged as most predictive, including terms such as “male,” “man,” “person,” “adult,” “talk,” and “look.” This further supports our finding that naturalistic stimuli with high social contents would elicit brain activity that has higher cognition predictive power. For sex prediction, the associated features were more abstract and general, including “entity,” “physical entity,” “object,” “living thing,” “artifact,” “whole,” and “causal agent” suggesting that sex-related neural patterns may be less tied to socially specific stimuli and more broadly distributed.

## 3. Discussion

In this study, we demonstrate the potential of deep neural networks (DNNs) to predict behavioral traits from the brain’s functional connectivity (FC) during movie watching, using data from the HCP 7T substudy. We show that while certain movie clips induce FC that has higher predictive power in terms of cognitive scores and sex than resting-state, others do not. We identify that movie clips with higher across-subject synchrony of brain activity also have better prediction performance for cognition, but not for sex classification. On a dynamic, finer temporal scale, we show that neural synchrony and prediction performance are significantly correlated over time for both cognition and, to a lesser extent, sex. Additionally, cognitive prediction accuracy is positively influenced by the presence of human faces and voices, while sex classification accuracy was not. Finally, we demonstrate that semantic content related to human and social stimuli enhances cognition prediction performance and content related to non-human objects enhances sex classification performance. We find that connections within and between sensory networks (VIS, SMN) and higher-order networks (DMN, FPN, SAN, DAN) are key for predicting both cognition and sex, with the most salient connections for both cognition and sex being similar but varying across different movie stimuli and rest. In conclusion, our findings highlight the importance of visual, auditory, and higher-order network connections in behavior prediction under naturalistic stimuli, underscore the role of neural synchrony in prediction accuracy, and suggest that human- and social-related movie content is especially effective in stimulating cognition-related brain activity.

The proposed deep neural network (DNN) for brain-behavior mapping introduces non-linearity and complex feature spaces, effectively capturing intricate relationships between functional connectivity (FC) and behavioral traits. We highlight that movie-watching FC yields a wide range of prediction accuracies for cognition and sex, depending on each clip’s ability to elicit neural synchrony and the clip’s content. Most clips outperform resting-state FCs of the same duration, and some even outperform rest sessions that are 5 times longer. While the model’s prediction results demonstrate its effectiveness, there are some limitations. Compared to previous studies on cognition prediction from movie-watching connectivity using linear models (connectome-based predictive modeling, CPM)[6], our results are similar overall, but the prediction accuracy is slightly higher for highly predictive clips (e.g. *Social Network*) and lower for less predictive clips (e.g. *Dreary*). We also note that some studies can achieve higher sex classification accuracy from a training set with more subjects and longer movie-watching [44]. This discrepancy may be due to overfitting, which can arise from the high complexity of the DNN, the high-dimensional input FC vectors, and the relatively small number of training subjects. To mitigate overfitting, we employed several strategies in the model design, applied strong regularization techniques, and reduced dimensionality via PCA. While the PCA+MLP framework performed better than other machine learning approaches (e.g., XGBoost, convolutional neural networks) we have tried, future research could explore more advanced non-linear approaches including topography-based networks [16], convolutional-recurrent models [19] and graph neural networks [45] [46].

Our findings also highlight the previously unexplored relationship between inter-subject neural synchrony during movie watching and FC prediction performance. There is a potential tradeoff in inducing neural synchrony - on one hand, higher synchrony improves the signal-to-noise ratio (SNR) of BOLD activity and corresponding FC; on the other hand, higher synchrony means less inter-subject variability, which may negatively impact behavior prediction. However, our overall finding of a positive correlation between neural synchrony and cognition prediction performance (and to a lesser extent sex classification) supports the conclusion that the primary benefit of synchrony lies in improving SNR of FCs - at least for this level of synchrony. While previous studies have pointed out that periods of high inter-subject FC similarity tend to align across viewers and are linked to the brain’s modular architecture [33], we here directly examine how similarity in BOLD responses affect brain-behavior mapping. Periods of higher synchrony tend to provide more accurate predictions, and brain regions of high synchrony tend to be more informative in terms of cognitive abilities and sex classification. These synchronizing moments are sometimes referred to as “events”, though in the context of co-fluctuations in z-scored BOLD activity across brain connections [29]. This finding is especially relevant for future fMRI experiment design, suggesting that stimuli that evoke higher synchrony may be preferable. It would be of interest to try and maximize synchrony to test if accuracy in predicting individual metrics would hit a ceiling - it is our opinion that such level of synchrony was not inducible due to inherent inter-individual variability in brain activity responses to the same stimuli.

At both static and dynamic scales, we found that the percentage of movie frames with human faces and voices was positively correlated with prediction accuracy for cognition, but not for sex. Additionally, we found that predictions of fluid and crystallized cognition are strongly associated with face presence (more so fluid), and to a lesser extent, voice presence, as shown in Supplementary Figure 2. These findings suggest that movies rich in social content engage brain networks relevant to cognition, but not to sex. This result aligns with previous findings at both the clip and dynamic levels: at the clip level, studies have shown that the duration of face presence is positively correlated with prediction accuracy during movie watching [6]; at the dynamic level, movie watching has been shown to elicit co-fluctuations in both core and extended face-processing regions, which in turn correlate with face recognition scores [47]. The stronger correlation between cognitive prediction and face presence than voice presence is also consistent with previous findings that face-selective brain regions show more robust and distributed identity-related neural patterns than voice-selective regions [48]. On the other hand, objects not related to humans - both living and non-living, were related to better sex classification. These findings have practical implications for designing future naturalistic viewing experiments — the content of the movie may be varied to obtain more accurate brain-behavior mappings, but this accuracy appears to depend on the specific outcome one is trying to predict.

Our implementation of DNNs in the brain-behavior mapping allowed us to employ the Integrated Gradients (IG) method to identify the most predictive connections on an individual subject basis. The gradient-based approach measures feature importance at the connection level, while the interpolation method enhances robustness. Our way to assess general feature importance across the population was to average the individual-level feature importances from the IG framework, however, one could also look at variability of the feature importances across individuals or subgroups within the overall group. We observe that the top FC features important for predicting cognition and sex are more consistent across movie clips than across the different outcomes that are being predicted, but that, generally speaking, the entire vector of feature importances were still moderately to highly correlated across movies (Fig. 3, 4, Supplementary Figure 4). The most important connections across the movie-watching FC mapping included connections within and between sensory networks (VIS, SMN) and higher-order networks (DMN, FPN, SAN, DAN). This finding is consistent with prior work using support vector machine models [25], which identified the FPN, DMN, and dorsal attention network (DAN) as key contributors to predicting cognitive states during movie-watching. It also partly agrees with work using convolutional models and deconvolution-based importance mapping [44], which showed that the default mode network (DMN) has the highest predictive power for sex classification, while the frontoparietal network (FPN) is most predictive of fluid and crystallized intelligence.

Importantly, our results reveal a high level of similarity between cognition and sex prediction importance maps within a clip compared to across a clip for the same outcome. Looking at the individual NIH Toolbox cognition test scores, we see a similar pattern of moderate levels of agreement across all movies (and rest) but tighter correlations within a movie for different cognitive scores than across a movie for the same cognitive score. This indicates that different movies may modulate somewhat different brain connections which could cause some differences in the feature importance maps. Interestingly, we see that the highest performing movie FC (Oceans 11, Social Network and Inception) feature importances are more similar to one another and tended to be more different from the other movie FC feature importance maps. Finally, the movie FC feature importances at both the regional and pairwise level were correlated with the FC feature importances from the resting-state model to an extent that was similar to the feature correlations across different movies. This may suggest that the neural substrates important for predicting to sex and cognition are similar across movies and rest, but that there is some variability induced by the specific movie being watched which likely depends on the across-subject synchrony of various brain regions and the resulting change in SNR of FC.

The analysis of movie-watching functional connectivity-based brain-behavior mapping is crucial for understanding the unique value of this paradigm in understanding the neural circuits underlying behavior traits or demographics. We find that two key elements correlated with movie-watching FC brain-behavior mapping accuracy are across-subject neural synchrony and movie features; we find that the movie features driving accuracy may depend on the metric we are trying to predict. Specifically, we find that movies with human- and social-related content are most effective for cognition and non-human object elements drive sex classification. In summary, these results highlight the importance of naturalistic viewing in fMRI neuroimaging and brain-behavior mapping, offering rich insights into brain mechanisms, and points us to further exploration of the mapping between movie features, neural synchrony, and accuracy in brain-behavior mapping using naturalistic paradigms.

## 4. Methods

### 4.1 Data

#### 4.1.1 Dataset, Participants, and Behavioral Measures

In this study, we use functional magnetic resonance imaging (fMRI) and behavioral data from the Human Connectome Project (HCP) 7T movie-watching dataset [41]. The dataset comprises n=184 healthy adult participants (ages 22 to 35), who underwent scanning using a Siemens Magnetom 7T scanner equipped with an optimized multiband echo-planar imaging (EPI) sequence (*TR* = 1000*ms, TE* = 22.2*ms, voxelsize* = 1.6*mm*^3^). We use a subset of n=176 subjects with complete data for all resting state and movie fMRI runs as well as behavioral data needed in this study.

In this study, we focus on predicting two items: biological sex (106 female, 70 male) and cognitive performance, both of which have been associated with functional connectivity patterns. We use the total cognition score which is the average of crystallized cognition and fluid cognition, which are derived from the NIH toolbox cognition tests including Picture Vocabulary, Reading Recognition, Dimensional Change Card Sort, Flanker Inhibitory Control and Attention, Picture Sequence Memory, List Sorting, and Pattern Comparison tests [49]. For completeness, we included the analysis for individual cognitive measures in Supplementary Information, but generally speaking the individual cognitive scores are noisier than the composite scores.

#### 4.1.2 fMRI Acquisition and Preprocessing

Resting-state fMRI (rfMRI) data were collected in four separate runs, each lasting approximately 16 minutes, at the beginning of each of the four 7T imaging sessions. During these runs, participants were instructed to keep their eyes open and maintain a relaxed fixation on a bright cross-hair projected against a dark background in a dimly lit scanning environment. For this study, we used data from the first (*REST1*) and fourth (*REST4*) resting-state sessions, as these were the only ones conducted on the same day as the movie-watching fMRI sessions.

Following the resting-state scans in the first and fourth imaging sessions, participants underwent two additional fMRI runs in which they viewed a total of 15 movie clips designed to engage cognitive and emotional processing. These clips, ranging from 1 to 4.3 minutes in length, consisted of both independent short films and excerpts from commercial Hollywood movies. The individual clips were concatenated into 12- to 14-minute-long .mp4 files, with a common Vimeo repeat validation clip included to assess test-retest reliability [50].

We excluded the test-retest clip, as well as another clip that was too short compared to the others. This resulted in a final set of 13 movie clips for our analysis. The movies were displayed using VisuaStim Digital MRI-compatible goggles to ensure consistent visual stimulation. We used brain activity data from all movie clips and resting-state sessions between the 10th and 174th seconds from the beginning of the scan (except where we also include all 15 mins of rsfMRI data for comparison), excluding the initial unstable period and aligning the end of the window with the shortest clip duration to ensure comparability of brain activity across all clips.

The specific movie clips analyzed in this study were as follows:

- MOVIE1 (first session, first day): Two Men, Welcome to Bridgeville, Pockets
- MOVIE2 (second session, first day): Inception, The Social Network, Ocean’s 11
- MOVIE3 (first session, second day): Flower, Hotel, Garden, Dreary
- MOVIE4 (second session, second day): Home Alone, Erin Brockovich, Star Wars

Clips in MOVIE1 and MOVIE3 are freely available independent films under Creative Commons licensing, and clips in MOVIE2 and MOVIE4 are from Hollywood movies as prepared and published by Cutting et al. 2012 [51].

Standard preprocessing of fMRI data, including motion correction, distortion correction, high-pass filtering, and nonlinear alignment to MNI template space [52] was used in the FIX-denoised volume-space data from the HCP 7T release. We also applied global signal regression (GSR) by removing the global mean time series signal from each voxel’s BOLD time series, thereby reducing potential confounds arising from widespread non-neuronal noise, including physiological artifacts and scanner-related fluctuations[53].

#### 4.1.3 Static and Dynamic Functional Connectivity Construction

##### Static functional connectivity(sFC)

To quantify static functional connectivity (sFC), we employed the Shen 268-region brain atlas [54]. For each region in the atlas, we computed the mean BOLD time series by averaging voxel-wise preprocessed signals within that region for each subject, run, and time frame. We then computed pairwise Pearson correlation coefficients between all 268 regions, yielding a 268 × 268 functional connectivity matrix for each movie clip or rest run per subject. We then extracted the upper triangular portion of each matrix (excluding the diagonal) and vectorized it into a 35,778-dimensional feature vector [6].

##### Dynamic functional connectivity(dFC)

To capture temporal fluctuations in functional connectivity during movie-watching, we adopted a sliding-window approach for dynamic functional connectivity (dFC). Each movie clip was segmented into overlapping time windows of 60 seconds, with a step size of 1 second, resulting in 104 sliding windows for each movie clip. Within each window, the mean BOLD signals were computed, and Pearson correlation was used to generate a 268 × 268 FC matrix for each window, resulting in 163 sliding-window dFC matrices for each clip.

### 4.2 Deep Neural Network(DNN) Prediction Model

We propose a novel deep neural network (DNN)-based behavior prediction model designed to capture the complex relationship between dynamic/static functional connectivity and behavioral and demographic traits (Fig. 1). The model takes as input a set of *n* = 176 functional connectivity vectors, each of size (35,778,), representing the upper triangular elements of the 268 × 268 FC matrix. The output corresponds to either cognition score (a continuous variable regression task) or biological sex (a binary classification task). Given the relatively small number of subjects, we employed a 10-fold cross-validation approach to ensure robust model evaluation. The 176 subjects were randomly divided into 10 groups, while making sure that individuals from the same family were always assigned to the same group. This step prevents data leakage and reduces the risk of inflated performance estimates due to genetic or familial similarities in brain connectivity. In each cross-validation iteration, the model was trained on 9 groups and tested on the remaining 1 group, ensuring that every subject was included in the test set exactly once. Predicted values for all subjects were collected across the 10 folds and then correlated with the true values to evaluate model performance. The division into 10 folds was performed 100 times to assess uncertainty due to varied groupings of individuals into train and test.

The prediction model consists of two primary components:

#### (1) Principal Component Analysis (PCA) Block

To reduce dimensionality and minimize noise, we apply PCA to the input FC vectors across all training subjects, retaining the first 100 principal components. This results in a (#training subjects, 100) feature matrix, capturing generally around 84% of the variance.

#### (2) Multi-Layer Perceptron (MLP) Block

- The PCA-reduced data is fed into a three-layer MLP, each layer containing 500 nodes with ReLU activation.
- To reduce overfitting, we employ dropout (rate = 0.5) in each hidden layer and apply L2 weight decay (0.01) as regularization.
- The model is optimized using the Adam optimizer with a learning rate of 0.001.
- The output layer is:
  – Linear for cognition prediction (regression task).
  – Sigmoid for sex classification (binary classification task).
- The loss function is:
  – Mean Squared Error (MSE) loss for cognition prediction.
  – Binary Cross-Entropy Loss for sex prediction.

While overfitting remains a challenge due to the limited sample size and the high-dimensional, noisy nature of fMRI-based FC measures [55] [56] [57] [58], the model’s hyperparameters were carefully tuned to strike a balance between learning the complex FC-behavior mapping and avoiding excessive overfitting. The model is developed using Pytorch on a Cuda GPU environment. To evaluate the model’s predictive performance, we computed Spearman’s correlation coefficient (*r*) between the predicted and true behavior scores for cognition. For sex classification, we assessed the model using the Area Under the Curve (AUC) of the receiver operating characteristic (ROC) curve.

### 4.3 Statistical Analysis of Predictive Connections and Networks

To quantify the predictive contribution of individual brain connections to the predictions, we employed Integrated Gradients (IG), a technique designed to estimate feature importance in deep neural networks [59] [60]. IG uses backward propagation to compute gradients of the model’s output with respect to the input, and improves robustness by integrating these gradients across a path between a baseline and the actual input.

In our DNN behavior prediction model, the IG procedure is as follows:

- PCA transformation: First, we perform Principal Component Analysis (PCA) on the training functional connectivity (FC) vectors, resulting in a (100,1) principal component(PC) vector for each training subject.
- Baseline Definition: The baseline reference input is defined as the average of each of the PC vectors across all training subjects (i.e. one vector per PC).
- Interpolation: A series of intermediate vectors are generated by linearly interpolating between the baseline and the actual input principle component (PC) vector of a subject. Specifically, for an input PC vector **x** and baseline 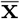, we compute the *k*th step:

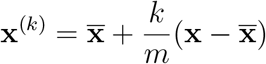

where *m* = 50 is the number of interpolation steps.
- Gradient Computation: For each interpolated vector **x**^(*k*)^, we compute the gradient of the DNN’s output (predicted cognition score or sex classification probability) with respect to its *i*th feature 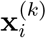, *i* = 0, …, 99.
- Integration Along the Path: The importance of each feature *i* of the PC vector is estimated by integrating these gradients over all interpolation steps:

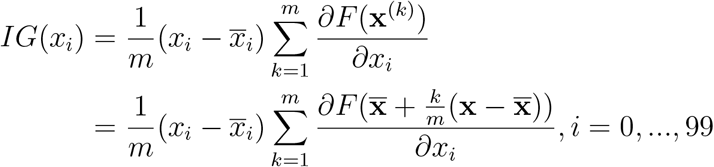

where *F* (*x*) is the DNN’s prediction function, and 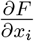 represents the gradient of the model output with respect to the *i*th entry of the input PC vector, computed by backward propagation.
- PCA inverse: The gradient is multiplied by the inverse of PCA loadings to project back to the gradients of the whole set of 35,778 brain connections.
- Feature Importance Assignment: For each subject, we obtain a set of IG values for each functional connection (i.e., each of the 35,778 connections). The absolute values of the IG for each connection are averaged across all subjects to compute its importance score, reflecting its contribution to behavior prediction.

At the regional level, we computed the average of the absolute values of the importance scores for the 267 connections associated with each region, which was used as its relative importance. At the functional network level, we mapped the most important connections to their corresponding Yeo canonical functional networks [42] and analyzed which functional networks contained the most predictive connections.

### 4.4 Neural Synchrony and its Relationship to Prediction Performance

#### 4.4.1 Neural Synchrony

During naturalistic viewing, people tend to exhibit higher neural synchrony when watching engaging and cognitively stimulating movies, whereas less engaging content often results in more variable neural responses across individuals [26] [27] [28] [29] [30]. To quantify neural synchrony, we calculated the inter-subject Pearson correlation of regional fMRI BOLD signals over time, reflecting shared neural responses across participants. This resulted in 268 region-specific synchrony values per subject pair. We then averaged these values across all subject pairs for each region to obtain regional population-level synchrony. For each movie clip, we identified significantly synchronized regions based on the Pearson correlation of BOLD signals across all subject pairs. To correct for the large number of pairwise comparisons (176×175*/*2 total pairs), we applied False Discovery Rate (FDR) correction using the Benjamini-Hochberg procedure [61]. Regions with adjusted p-values below *α* = 0.05 were considered significantly synchronized. We took the union of significant regions across clips (collected into the set called ROI), and then averaged these to obtain a whole-brain synchrony measure; this procedure excludes regions with non-significant synchrony across all movie clips.

- Pairwise neural synchrony for a pair of subjects (*i, j*), for a region *v*:

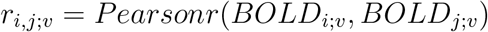
- Region-wise synchrony for a region *v* for all subject pairs:

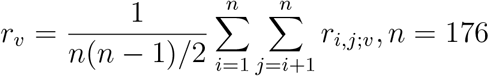
- Global, whole brain synchrony for a pair of subjects (*i, j*):

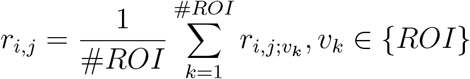

#### 4.4.2 Relationship Between Neural Synchrony and Behavior Prediction Performance

To investigate the relationship between neural synchrony and behavior prediction, we analyzed both whole-brain and regional levels. At the whole-brain level, we computed an overall synchrony score for each movie clip by averaging synchrony across all regions of interest (ROIs). We then correlated this score with the prediction accuracy for cognition and sex across all movie clips to assess whether a significant relationship exists.

At the regional level, we examined the correlation between the region-wise synchrony map and the region-wise prediction importance map. The prediction importance map was calculated by averaging the importance scores of all 267 functional connections linked to each brain region. A positive correlation would indicate that regions with higher synchrony are also more predictive. To verify the robustness of the correlation, we used both Pearson and Spearman correlation metrics. This dual approach helps ensure the correlation is not driven solely by low-synchrony regions, where predictive power might still vary despite minimal synchrony.

To further evaluate the impact of high- and low-synchrony brain regions on behavior prediction, we conducted targeted predictions using subsets of brain regions exhibiting distinct levels of synchrony. For each movie clip, we identified high-synchrony regions as those with inter-subject correlation values satisfying FDR-corrected p<0.05. We then selected an equal number of randomly chosen regions and an equal number of low-synchrony regions (ranked in descending order of p-values). Using these subsets, we trained separate DNN prediction models while maintaining the same model architecture and compared their prediction accuracies. This approach allowed us to determine whether high-synchrony regions contribute more strongly to behavior prediction compared to randomly selected or low-synchrony regions.

### 4.5 Sliding-Window Dynamic Functional Connectivity, Neural Synchrony and Movie Features

#### 4.5.1 Sliding-Window Dynamic Functional Connectivity

To capture temporal dynamics, we employed a sliding-window approach that segments each 164-second movie clip into overlapping 60-second windows with a 1-second step size. This resulted in 104 windows per movie clip, with each window defined by its midpoint (e.g., the 30th second corresponds to the window from 0 to 60 seconds).

We trained separate DNN models to predict cognitive score and sex for each 60-second sliding window dynamic functional connectivity (dFC), ensuring that the model performance reflected the predictive power of the FC for that specific time window under the given naturalistic stimulus. While the structure of the DNN model and cross-validation method were identical to those used for static FC, the key difference was that the input FC vector was based on shorter temporal segments. This approach allowed us to generate an accuracy curve over time during the movie clip, which we later compared with fluctuations in neural synchrony and movie features.

#### 4.5.2 Sliding-Window Neural Synchrony

We use the same set of overlapping 60-second sliding windows and compute the intersubject correlation of fMRI BOLD time series within the window to generate a dynamic sequence of whole-brain neural synchrony levels. Similar to static neural synchrony, we took the average of inter-subject correlation across regions of interest (ROIs) to obtain the whole-brain synchrony level for each window. We can thus obtain the time fluctuation of synchrony level among the subjects. At the movie clip level, we computed the Pearson correlation between neural synchrony and dynamic FC prediction accuracies across the 104 windows for each clip. A strong positive correlation would indicate a high degree of co-fluctuation and similar temporal patterns between synchrony and FC prediction performance over time. At a more aggregate level, we combined the sliding windows from all 13 movie clips and computed the same Pearson correlation between neural synchrony and dynamic FC prediction accuracies across the 13×104 windows. This approach provides much more data points compared to static FC, allowing for a more robust analysis and helping to further validate the relationship between synchrony and behavioral prediction performance.

#### 4.5.3 Sliding-Window Movie Features

We also investigated whether specific high-level features of the movie, including the presence of human faces and voices, influenced the accuracy of behavior prediction. To quantify these features, we calculated the percentage of time each feature was present within each 60-second sliding window, defined as the proportion of TRs (time points) where at least one human face or one person talking was detected. The presence of human faces in each frame was identified using the open-source Python package *pliers* [62]. Human voice presence in each frame is identified using the open-source python package *webrtcvad* (https://github.com/wiseman/py-webrtcvad). Note that the TR for fMRI scanning in the HCP-7T dataset is 1 second, meaning the “frame” here is also 1 second to align with the dataset’s temporal resolution. We then examined the correlations between these features and the sliding-window dFC (dynamic functional connectivity) prediction accuracy to explore how visual and auditory elements shape dFCs predictive power.

Additionally, we used WordNet features to analyze the semantic content driving high prediction accuracy for cognition and sex. The HCP-7T dataset includes frame-wise word descriptions that label the content of each frame, as specified by the HCP movie labeling pipeline [63] [64]. For each sliding window, we counted the occurrences of each word, then weighted these occurrences by the behavior prediction accuracy for that window. The average of these weighted occurrences was computed across all 13×104 sliding windows. This produced a weighted average for each word in the movie labeling dictionary, reflecting the words most influential for behavior prediction which we visualized using word clouds.

## 6. Data Availability

The HCP 7T Dataset is publicly available at https://db.humanconnectome.org/. Preprocessed shen268 atlas movie-watching BOLD time series data can be found at: https://github.com/esfinn/movie_cpm.

## 7. Code Availability

The code used in this study, along with usage instructions, is publicly available at: https://github.com/Kyrrego/MovieFC_Behav_Mapping.git.

## 8. Acknowledgements

This work was supported by the Ann S. Bowers foundation through the Ann S. Bowers Women’s Brain Health Initiative.

## 10. Author Contributions

A.K. and M.S. conceived the experiments and interpreted the results. Y.Z. conducted the experiments, analyzed, and interpreted the results. E.F. interpreted the results. Y.Z. and A.K. wrote the manuscript. All authors reviewed the manuscript.

## Appendix

Please see supplementary information in additional files.

