## Supplementary Figures for "View, engage, predict: enhancing brain-behavior mapping with naturalistic movie-watching fMRI"

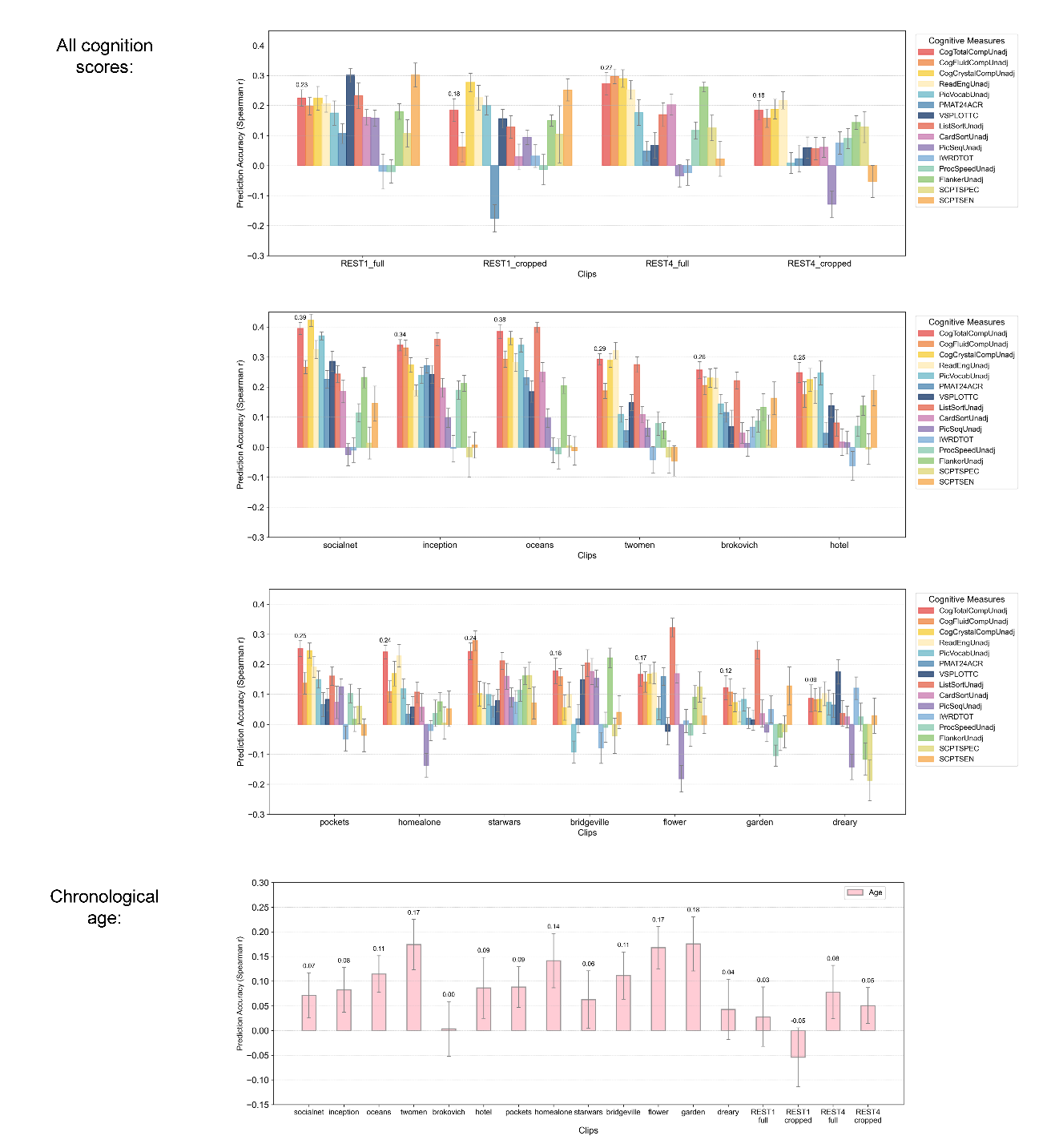

Supplementary Figure 1: Supplement of Fig.2. Prediction accuracies for all cognitive measures from the NIH Toolbox and chronological age, computed for each movie clip from the HCP 7T release. Here, we show results of both the first 164 seconds of the rest sessions and the 15-min full rest sessions.

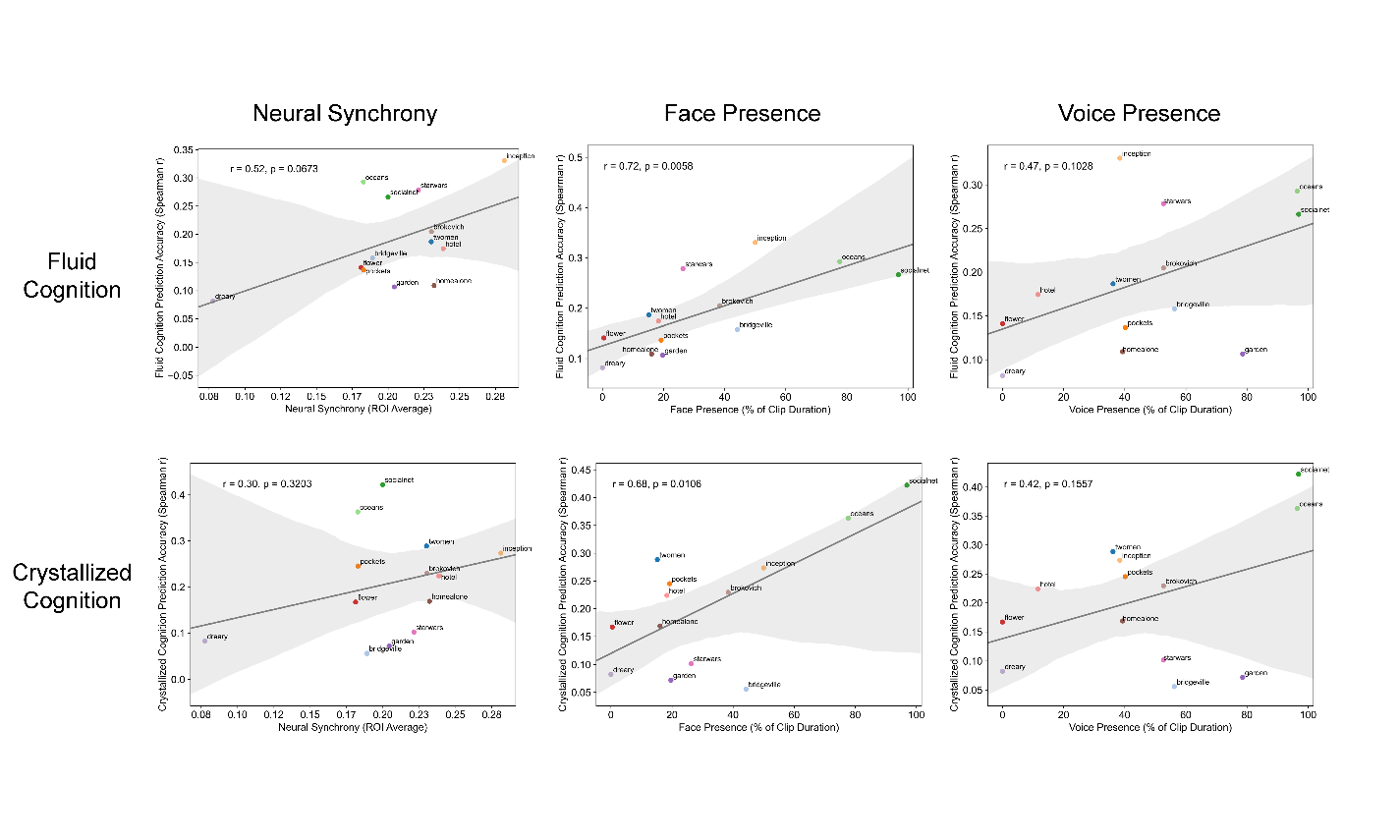

Supplementary Figure 2: Supplement of Fig.2. Correlation between movie clip FC-based prediction accuracy for fluid/crystallized cognition and neural synchrony, face and voice presence duration.

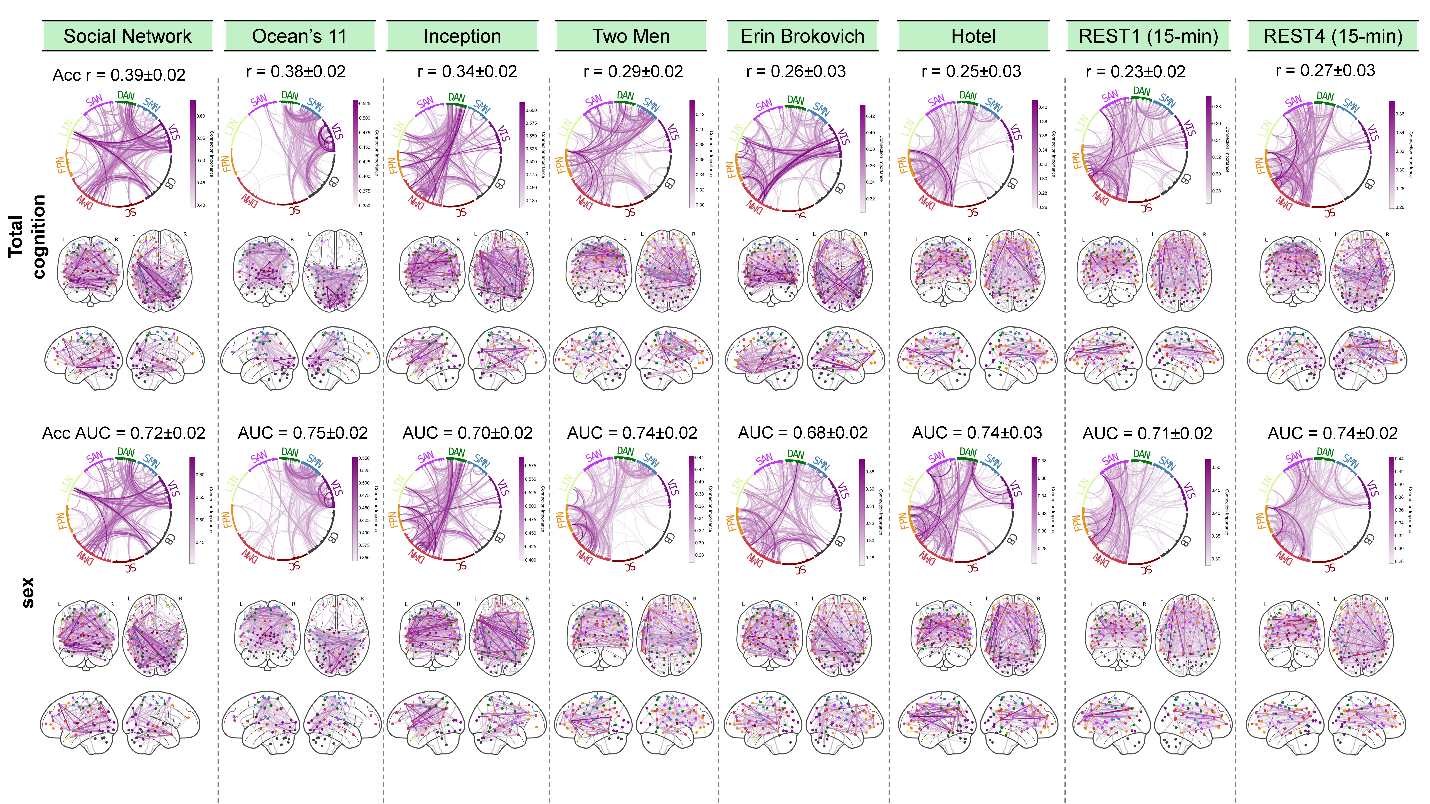

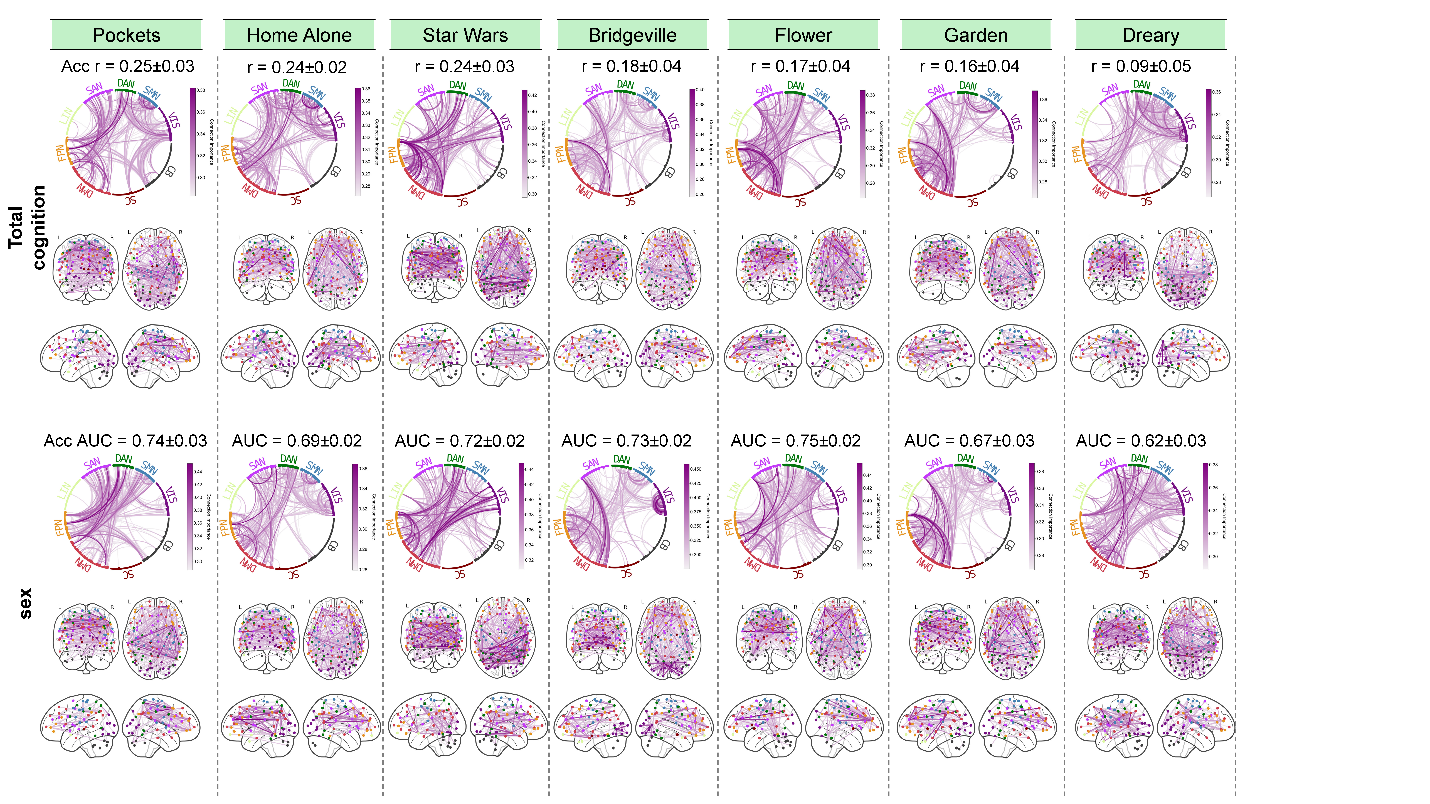

Supplementary Figure 3: Supplement of Fig.3. For each clip and resting-state session, chord and corresponding glass brain diagrams visualize the most important connections—specifically, those within the top 1% of average importance scores—for both total cognition and sex prediction.

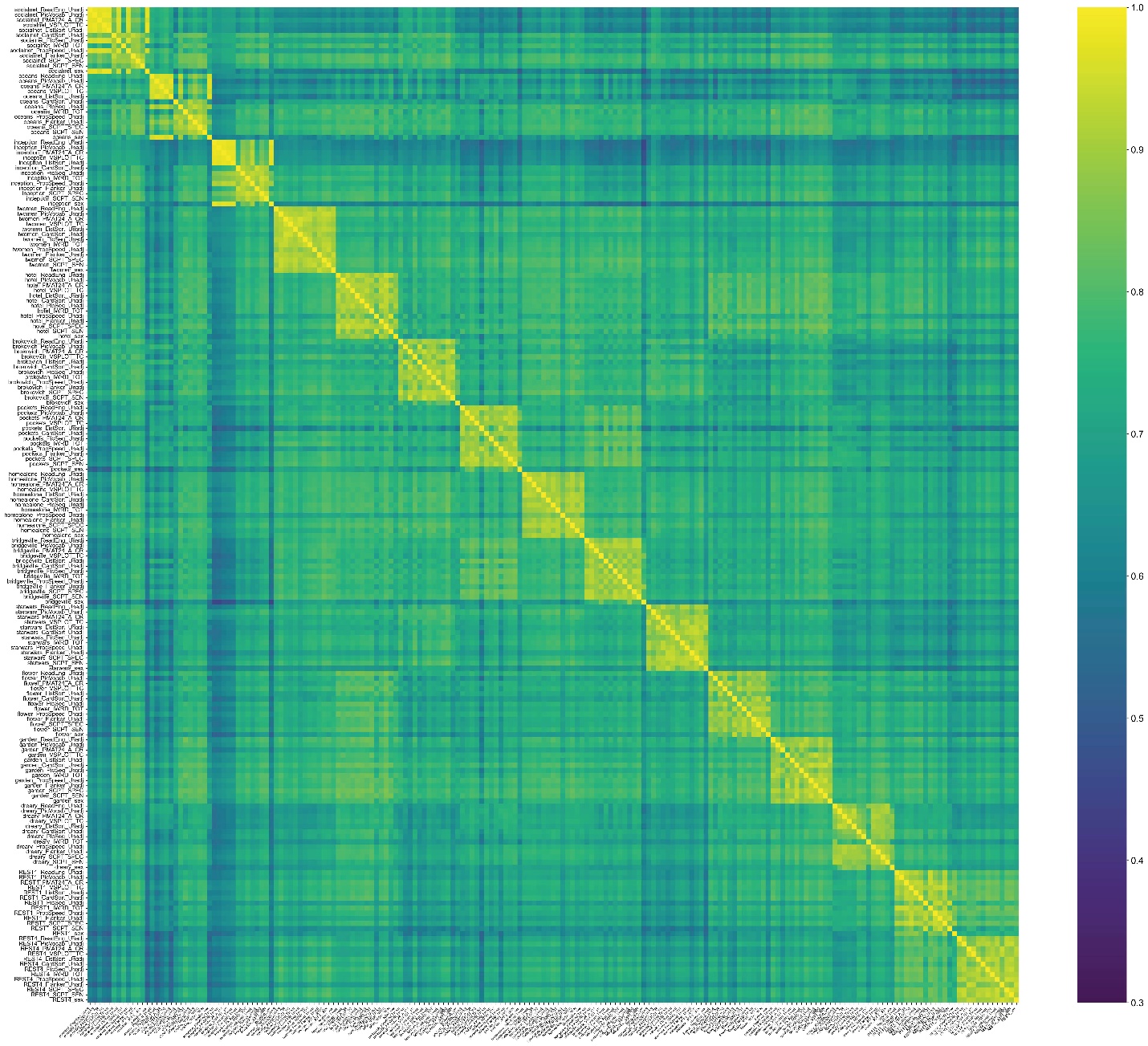

Supplementary Figure 4: Supplement of Fig.3. Correlation of connection-level prediction importance maps for all movie clips and all 12 individual NIH toolbox cognition scores, plus sex. For some movie clips, the importance maps for sex and some cognition scores are less correlated than others (e.g. for *Social Network*, *Ocean's 11* and *Inception*, the Dimensional Change Card Sort Test score shows lower correlation to other cognitive scores). However, the higher correlation values near the diagonal show that importance maps tend to be more similar within a given clip for the varied cognition and sex predictions as compared to the same cognitive score/sex across different clips.

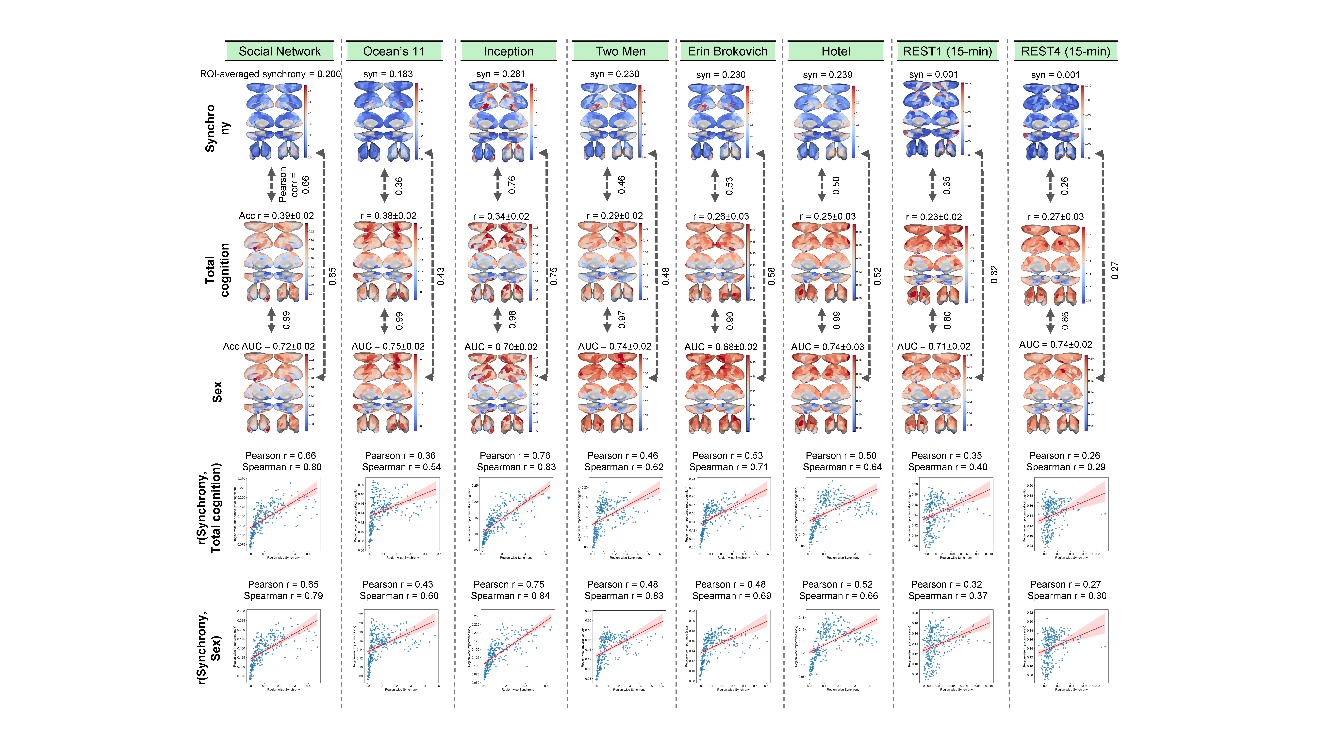

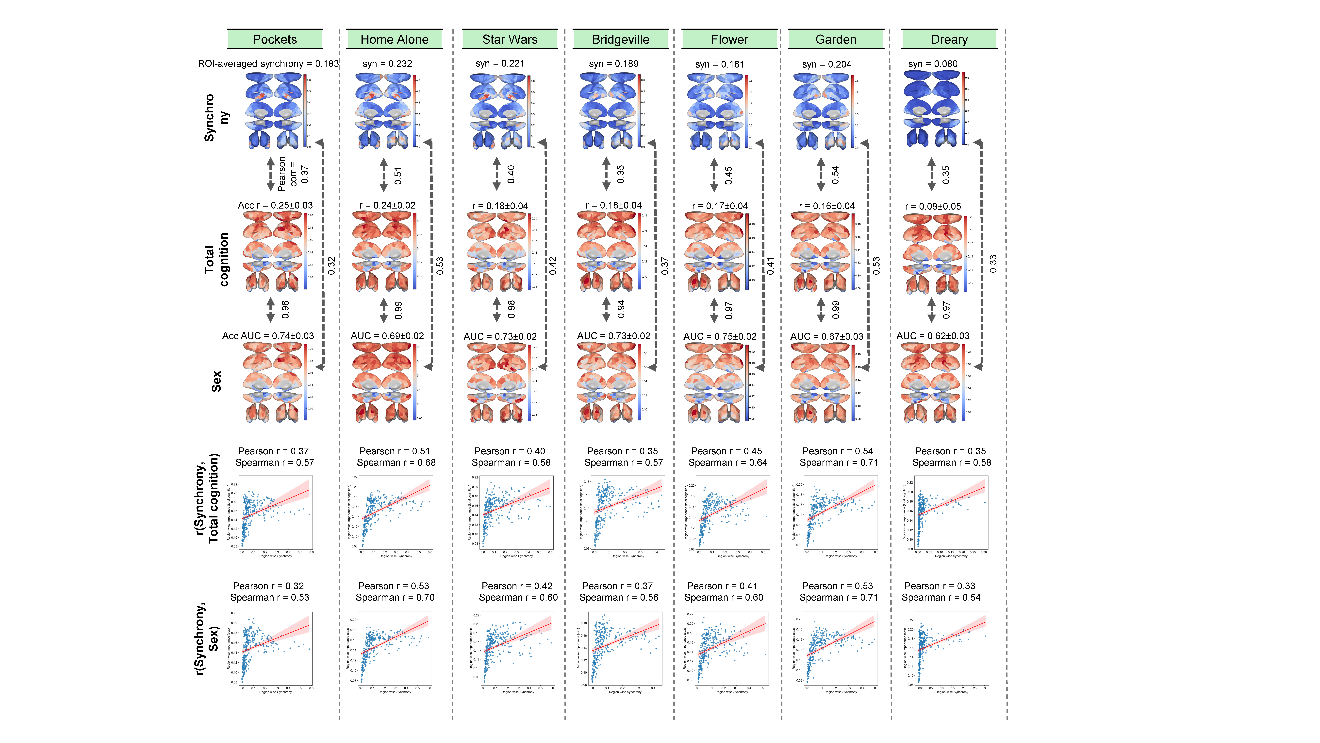

Supplementary Figure 5: Supplement of Fig.4. For each clip, the first row shows region-wise neural synchrony maps. The second and third rows display region-wise prediction importance maps for total cognition and sex. The importance of each region is calculated as the average of the importance scores of all connections connected to that region. The fourth and fifth rows show the Pearson correlation between region-wise neural synchrony and prediction importance maps for total cognition and sex.

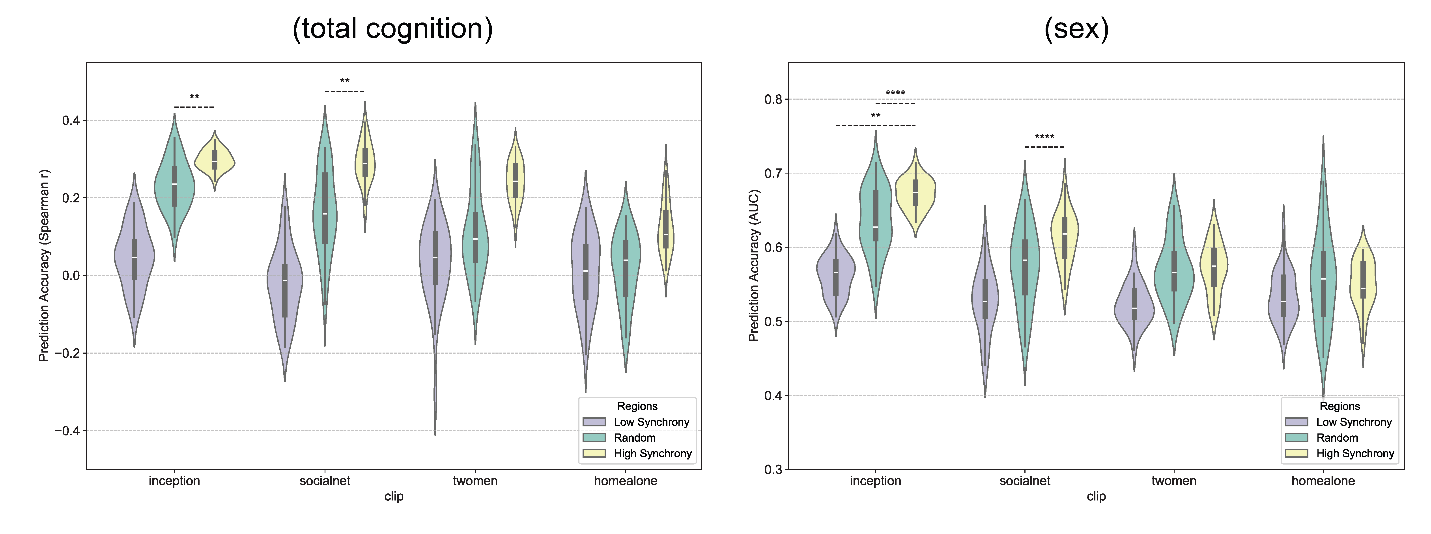

Supplementary Figure 6: Supplement of Fig.4. Prediction performance using an equal number of high-synchrony, low-synchrony, and randomly selected regions. For both cognition and sex prediction, models using FC of high-synchrony regions outperform models based on random regions' FC, which in turn outperform models based on FC from low-synchrony regions.
